# Histone 3 lysine 36 trimethylation by SETD2 shapes an epigenetic landscape in intestinal stem cells to orchestrate lipid metabolism and prevent cell senescence

**DOI:** 10.1101/2025.04.09.648070

**Authors:** Yue Xu, Ziyi Wang, Wenxin Feng, Hanyu Rao, Dehuan Wang, Wei Zhang, Rebiguli Aji, Ningyuan Liu, Wei-Qiang Gao, Li Li

**Author notes:** Equal contribution (Yue Xu, Ziyi Wang).

## Abstract

The self-renewal capacity of intestinal stem cells (ISCs) declines with aging, leading to a loss of homeostasis and an increased susceptibility to intestinal diseases. Despite the established significance of lipid metabolism and epigenetic regulation in ISC function, the molecular mechanisms that connect these processes to aging-related ISC dysfunction remain elusive. Here, we demonstrate that Histone 3 lysine 36 trimethylation (H3K36me3) caused by SETD2 is critical for ISC stemness. We found that H3K36me3 deficiency results in reduced ISC proliferation and differentiation, disrupts fatty acid oxidation (FAO) metabolism, and induces ISC senescence. Mechanistically, the loss of H3K36me3 triggers the activity of the SWI/SNF chromatin remodeling complex and leads to increased chromatin accessibility and enhancer activation, which alters FAO- and senescence-related gene expression. Importantly, we discover that metabolic intervention can prevent the senescence of ISC due to H3K36me3 deficiency. Our findings reveal a crucial role for H3K36me3 in maintaining the epigenetic landscape that orchestrates FAO metabolism and determines intestinal stem cell functions, emphasizing the role of FAO as a key modulator between H3K36me3 and ISC aging, suggesting that metabolic intervention may help mitigate age-related ISC dysfunction.

## Introduction

The intestinal epithelium is one of the most rapidly self-renewing tissues in adult mammals. Intestinal stem cells (ISCs), located at the base of the crypts, play a crucial role in villus formation and epithelial integrity by sustaining a high rate of self-renewal. The self-renewal capacity and differentiation potential of ISCs decline with aging, leading to a loss of intestinal epithelium homeostasis^1–3^. This decline in stemness accelerates the aging process of ISCs, promoting cellular senescence and increasing susceptibility to intestinal diseases^4–8^. Given the high energy demands associated with rapid cell division and epithelial renewal, ISCs are highly dependent on fatty acid oxidation (FAO) ^9–13^. Multi-omics analyses have demonstrated that the expression of FAO-associated transporters and enzymes declines more rapidly with age^14^, suggesting a connection between aging and metabolic alterations in ISCs. Fasting has been shown to enhance ISC function in aged mice by activating FAO^15^, underscoring the critical role of lipid metabolism in ISC function, particularly during aging. However, the molecular mechanisms underlying age-related ISC dysfunction remain complex and are not yet fully elucidated.

Histone modification plays a pivotal role in determining ISC fate^16–18^. Histone methylation, characterized by numerous methylation sites and differentially methylated states, regulates gene expression by modulating chromatin accessibility, thereby influencing the transcriptional programs that govern ISC function^19–22^. Among these modifications, H3K36 methylation is particularly important for maintaining genomic stability, with SETD2 serving as key enzymes responsible for the trimethylation of H3K36^23^. Deficiencies of SETD2 can lead to developmental defects and disease^24,25^. It has been reported that a loss of H3K36me3 results in a more open chromatin conformation, potentially increasing cryptic transcription during aging^26^. Recent studies have further revealed that H3K36 methylation regulates cell plasticity and regeneration in the intestinal epithelium^27^. However, the specific role of H3K36me3 in ISC senescence remains poorly understood.

Here, we demonstrate that H3K36me3, caused by SETD2, is critical for maintaining ISC stemness. We find that loss of H3K36me3 leads to reduced proliferation and differentiation of ISCs, disrupted lipid metabolism, and induced senescence. Specifically, the loss of H3K36me3 alters the function of the SWI/SNF chromatin remodeling complex, leads to increased chromatin accessibility and enhancer activation to affect gene transcription. Notably, metabolic interventions can rescue the decline in ISC numbers caused by H3K36me3 loss. These findings shed light on the role of H3K36me3 in ISC biology and may inform strategies for maintaining intestinal homeostasis and developing treatments for age-related intestinal disorders and stem cell dysfunction.

## Results

### The levels of H3K36me3 are reduced in the intestinal stem cells of aged mice

To study whether H3K36me3 changes during the development of ISCs, we analyzed the small intestinal epithelium of Lgr5-eGFP-IRES-CreERT2 reporter mice^28^, as Lgr5 is a well-established marker of ISCs. As expected, we observed that aging led to an increase in SA-β-galactosidase (SABG)-positive cells (a widely recognized cell senescence marker^29^). (Fig. 1A-B). Immunofluorescent staining analysis demonstrated that H3K36me1 and H3K36me2 were highly expressed in ISCs at the early developmental stages but reduced in aged mice (Fig. 1A and 1C). In contrast to H3K36me1 and H3K36me2, H3K36me3 exhibited a distinct expression pattern. The levels of H3K36me3 were lower at P0, 1 week, 3 weeks, and 5 weeks but gradually increased from 7 weeks, peaking at 2 months (Fig. 1A and 1C). Furthermore, H3K36me3 was also significantly reduced with advanced age, particularly in 20-month-old mice (Fig. 1A and 1C), suggesting a potential association between H3K36me3 and ISC senescence.

**Figure 1.**
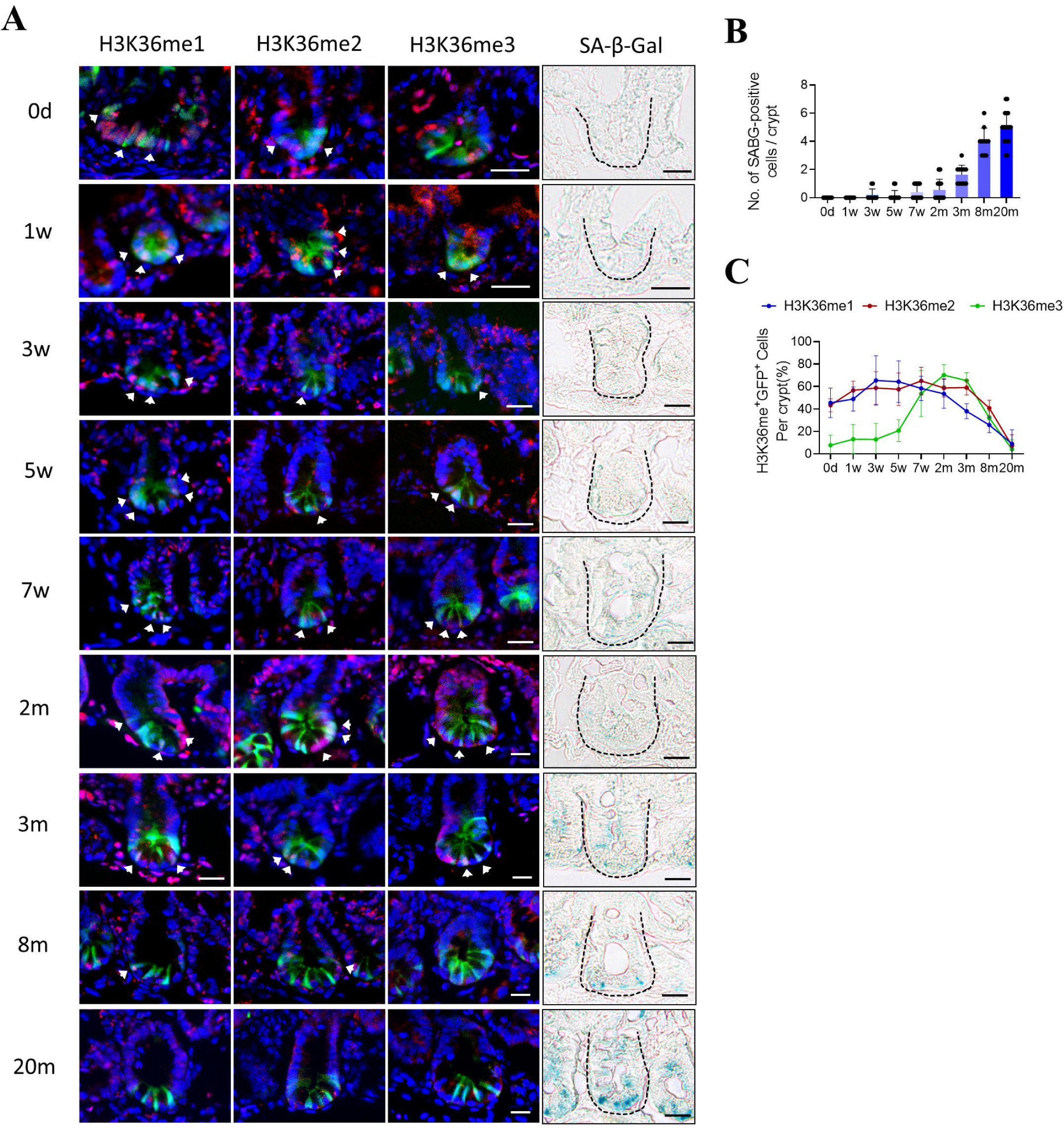
Histone 3 lysine 36 methylation levels are reduced in the intestinal stem cells of aging mice. **A.** Frozen sections of various timepoint mouse intestine tissues were stained with indicated histone modifications (red) or SA-β-Gal (blue) (n = 3). Scale bars, 20 μm. **B.** Quantifications of numbers of SA-β-Gal positive senescent cells per crypts were measured. **C.** Quantifications of indicated histone modifications (red) in Lgr5-EGFP+ ISCs per crypts were measured.

### H3K36me3 loss impairs intestinal stem cell proliferation and differentiation

Given the observed alterations in H3K36me3 levels across different developmental stages of ISCs and its correlation with ISC senescence, we sought to investigate the impact of H3K36me3 deficiency on the stemness of ISCs. We specifically deleted SETD2 in *Lgr5-EGFP-CreERT2;Setd2*^f/f^ (*Setd2*^ISC-KO^) mice. After five days of tamoxifen induction, *Setd2* expression was lost (Extended Data Fig. 1A), and H3K36me3 was essentially absent from the ISCs in the knockout mice (Extended Data Fig. 1B). We then analyzed the number of Lgr5-GFP^+^ cells following tamoxifen treatment (Fig. 2A). Our results indicated that the deletion of *Setd2* resulted in a decreased number of Lgr5-GFP^+^ cells after tamoxifen treatment (Fig. 2B-C and Extended Data Fig. 1C). Additionally, the transcription levels of Lgr5 stem cell markers, including *Lgr5*, *Olfm4*, *Ascl2*, *Sox9*, *Lrig1*, *Tert* and *Bmi1*, were significantly downregulated (Fig. 2D). These findings suggest that H3K36me3 is essential for the stemness maintenance of ISCs.

**Figure 2.**
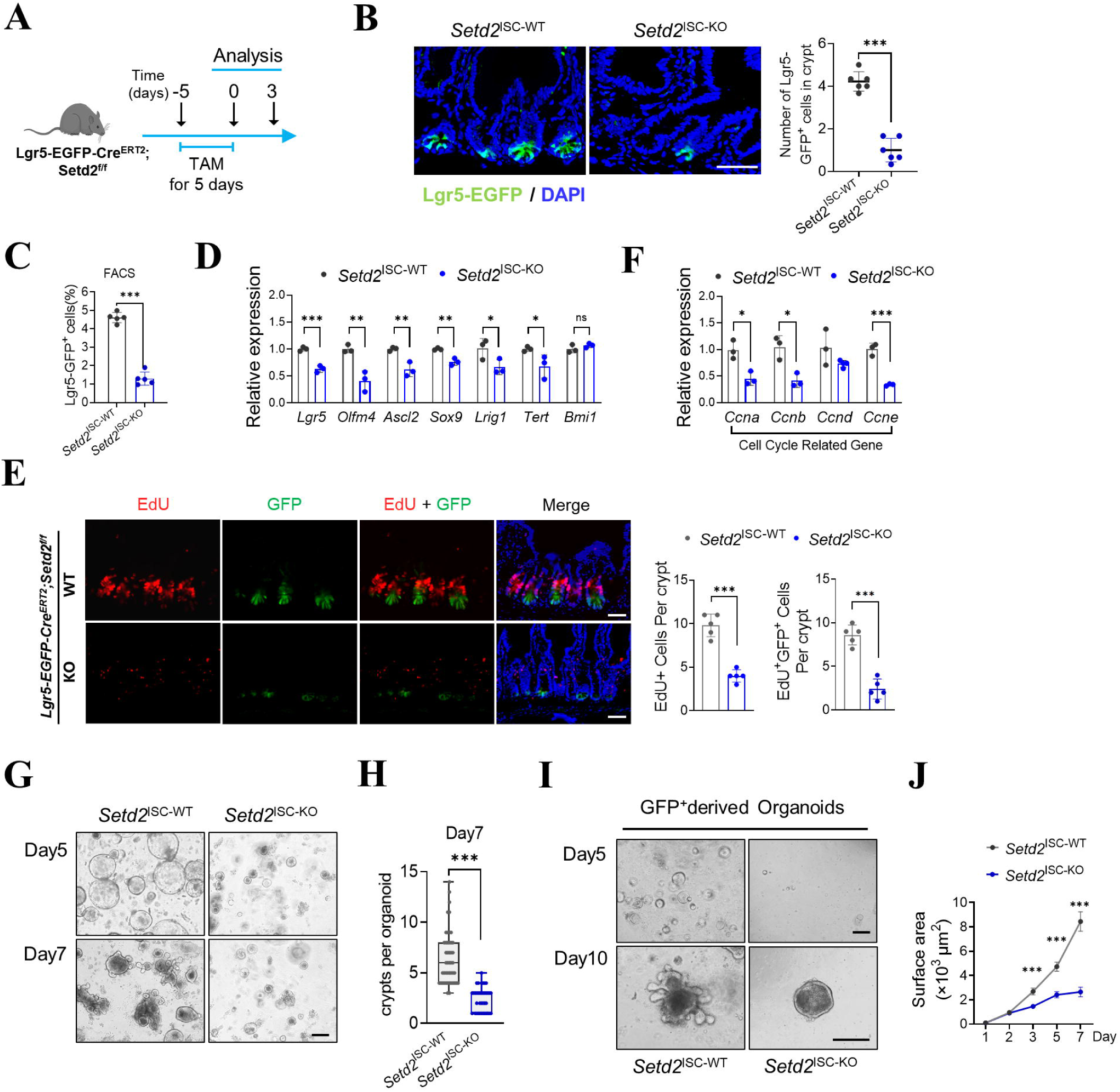
H3K36me3 loss impairs intestinal stem cell proliferation and differentiation. **A.** A schematic for conditional ablation of SETD2 by tamoxifen treatment in *Lgr5-EGFP-CreERT2; Setd2*^f/f^, respectively. **B.** Representative graphs of crypts containing Lgr5-EGFP+ ISCs in the intestinal sections after tamoxifen treatment. Scale bars, 50 μm. **C.** Determination of Lgr5–EGFP^hi^ cells from *Setd2*^ISC-WT^ and *Setd2*^ISC-KO^ mice by flow cytometry, n = 5 mice per group. **D.** Expression of stem cell markers in ISCs isolated from the small intestine. **E.** Representative images and quantification of EdU incorporation after a 2-h EdU (red) pulse in the intestinal sections from *Setd2ISC-WT* and *Setd2*^ISC-KO^ mice. Scale bar, 50 µm, n = 3 mice per group. Data are mean ± s.e.m. of biologically independent samples. Statistical significance was determined by a two-tailed Student’s t-test. **F.** Expression of cell cycle-related genes in ISCs isolated from the small intestine. **G.** Crypt number of organoids derived from isolated crypts. Data was pooled from 3 independent experiments with n ≥ 3 mice per group. **H.** Representative images of organoids formed by crypts from *Setd2*^ISC-WT^ and *Setd2*^ISC-KO^ mice on day 5 and day 7. Scale bars, 200 μm. **I.** 10000 Lgr5-EGFP^high^ ISCs were sorted from the small intestine of *Setd2*^ISC-WT^ and *Setd2*^ISC-KO^ mice. Representative images of organoids are shown from two independent experiments, Scale bars, 200 μm. **J.** Quantification of the surface area of organoids formed by crypts as described in **I.** Data was pooled from 2 independent experiments, >30 organoids were analyzed for each group. For qRT-PCR, GAPDH was used as an internal control. The statistical data represent mean ± s.d. (n = 3 mice per genotype). Student’s t-test: ns, non-sense. *P < 0.05. **P < 0.01. ***P < 0.001. All images are representative of n = 3 mice per genotype.

To evaluate the effects of H3K36me3 loss on ISC proliferation in vivo, we quantified proliferating cells in the small intestine of tamoxifen-treated *Setd2*^ISC-KO^ mice using an EdU assay. As shown in Fig. 2E, the number of EdU-positive cells was significantly lower in *Setd2*^ISC-KO^ mice, which was further supported by a marked reduction in the S/G2/M phase cell population (Extended Data Fig. 2D). Consistent with these results, the expression of genes that promote the cell cycle was diminished in the epithelium of *Setd2*^ISC-KO^ mice (Fig. 2F). Additionally, we observed that the deficiency of H3K36me3 in ISCs significantly influenced their differentiation into various epithelial cell types, including goblet cells, tuft cells, and Paneth cells (Extended Data Fig. 2E-H). Collectively, these results highlight the crucial role of H3K36me3 in both the proliferation and differentiation of Lgr5^+^ stem cells.

In order to further elucidate how H3K36me3 loss interferes with ISCs function, we tested the potential of isolated intestinal crypts to form clonal, multipotent organoid bodies in the absence of SETD2. Remarkably, crypts isolated from *Setd2*^ISC-KO^ mice failed to self-expand into organoids and exhibited less budding compared to control mice (Fig. 2G-H). To investigate the cell-autonomous role of H3K36me3 in Lgr5^+^ ISCs, we isolated Lgr5^+^ cells from the small intestines of control or *Setd2*^ISC-KO^ mice. After 5 days culture in conditioned growth medium, the isolated Lgr5^+^ cells expanded and self-organized into spherical organoids. However, the organoid spheroids derived from *Setd2*^ISC-KO^ mice were significantly smaller in size and had fewer crypts compared to those from controls (Fig. 2I-J), indicating that *Setd2*-deficiency ISCs were much less efficient at generating organoids. Collectively, our findings demonstrate that H3K36me3 is indispensable for ISC function under homeostatic conditions.

### H3K36me3 in ISCs disturbs lipid metabolism and leads to senescence

Next, to further investigate the mechanism of H3K36me3 in ISCs, we performed RNA sequencing (RNA-seq) on Lgr5^+^ISCs (EpCam^hi^, CD24^low^, EGFP^hi^) sorted from both *Setd2*^ISC-WT^ and *Setd2*^ISC-KO^ intestinal tissue, respectively (Fig. 3A). The data revealed significant changes in gene expression in the *Setd2*^ISC-KO^ mice, with 1,337 genes upregulated and 2,831 genes downregulated (log2(fold change) > 0.585, p < 0.05) (Fig. 3B and Extended Data Fig. 2A). GSEA analysis indicated that DNA replication and cell cycle progression were significantly downregulated upon loss of H3K36me3 (Fig. 3C and Supplementary Table 1). GO analysis showed that the most prominent alteration was an enrichment in lipid metabolism process, including fatty acid oxidation and lipid transport (Fig. 3D and Extended Data Fig. 2B), as confirmed by RT-qPCR analysis (Extended Data Fig. 2C). Lipid quantification and fluorescence staining of intestinal crypts confirmed that *Setd2*^ISC-KO^ mice exhibited increased lipid accumulation (Fig. 3E-F).

**Figure 3.**
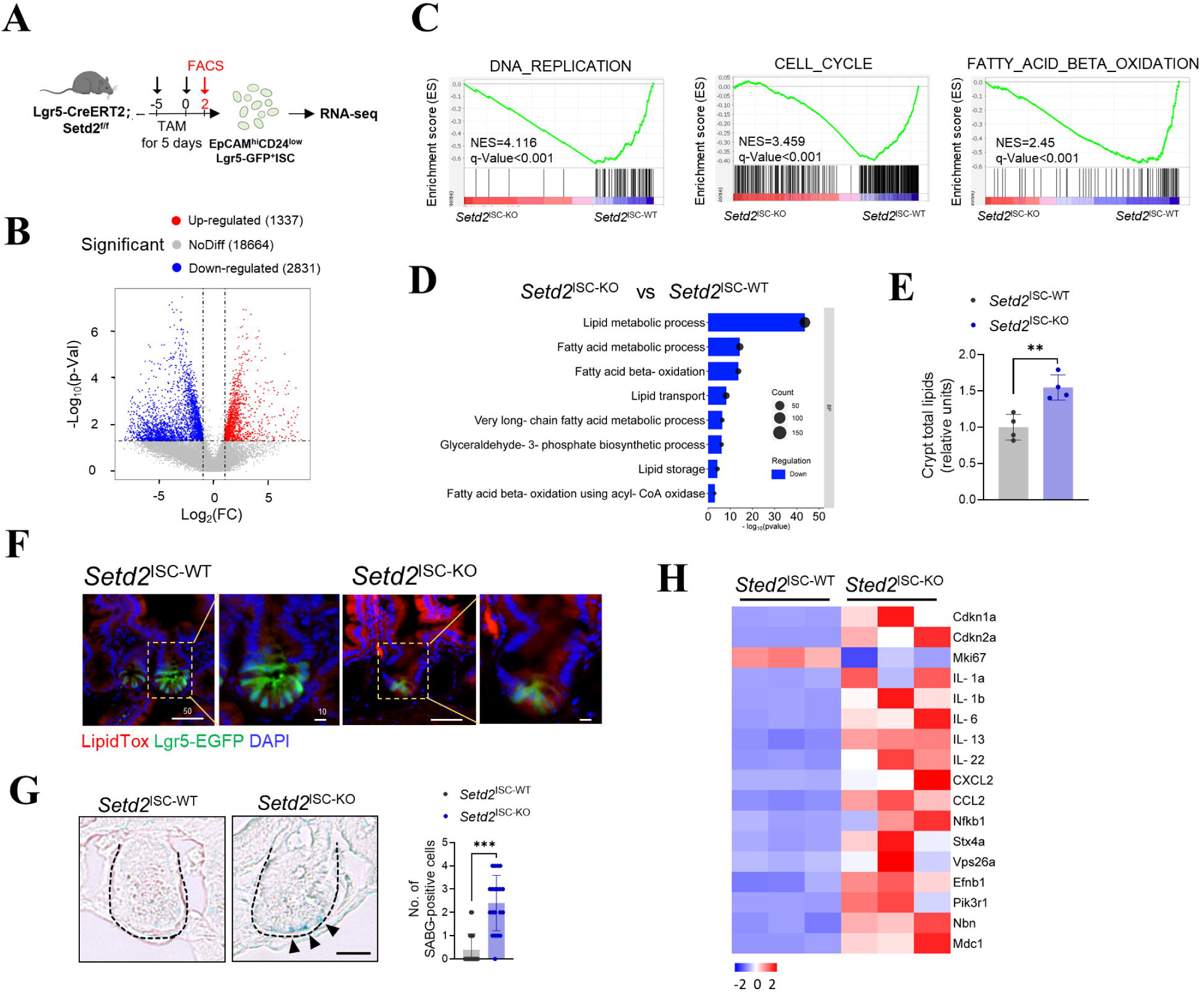
H3K36me3 loss in ISCs disturbs lipid metabolism and leads to senescence. **A.** Schematic for conditional ablation of SETD2 by tamoxifen treatment. **B.** Volcano plots displaying differentially expressed between *Setd2*^ISC-WT^ and *Setd2*^ISC-KO^ mice (P < 0.05, two-sided Student’s t-test). Red and blue dots represent upregulated and downregulated genes, respectively. **C.** GSEA of RNA-seq in *Setd2*^ISC-KO^ mice compared to *Setd2*^ISC-WT^. NES, normalized enrichment score. **D.** RNA-seq of the intestines from *Setd2*^ISC-WT^ and *Setd2*^ISC-KO^ mice was conducted. Gene Ontology (GO) pathway analysis identified lipid metabolism-related pathways affected by the deletion of SETD2 in ISCs. **E.** Relative total lipid concentrations in isolated IECs from *Setd2*^ISC-WT^ and *Setd2*^ISC-KO^ mice. **F.** LipidTox detection of fatty acids in the small intestines of *Setd2*^ISC-WT^ and *Setd2*^ISC-KO^ mice. Scale bars, 50 μm (zoom, 10 μm). **G.** Representative images showed decreased SA-β-Gal positive senescent cells from *Setd2*^ISC-WT^ and *Setd2*^ISC-KO^ (n = 3). Scale bar, 20 μm. **H.** Heatmap visualizing expression levels of selected cell senescence-related genes with altered expression in the ISCs of *Setd2*^ISC-WT^ and *Setd2*^ISC-KO^ mice. For qRT-PCR, GAPDH was used as an internal control. The statistical data represent mean ± s.d. (n = 3 mice per genotype). Student’s t-test: *P < 0.05. **P < 0.01. ***P < 0.001. All images are representative of n = 3 mice per genotype.

We further conducted a metabolomic analysis on crypts from *Setd2*^ISC-KO^ mice. Principal component analysis (PCA) of the metabolomic data revealed a distinct separation between the metabolite profiles of ISCs with H3K36me3 deficiency and those of control ISCs (Extended Data Fig. 3B). Using a false discovery rate (FDR)-corrected Wilcoxon test, we identified 102 differentially abundant (DA) metabolites in the intestinal crypts of *Setd2*^ISC-KO^, compared to controls (FDR-adjusted p < 0.05) (Extended Data Fig. 3C). Among these metabolites, we focused on fatty acids (FAs) with 20 or more carbon atoms, which are selectively metabolized by β-oxidation in peroxisomes (Extended Data Fig. 3D). Consistent with our observations in Figure 3, the quantified free fatty acids (FFAs) showed an increase in *Setd2*^ISC-KO^ mice, suggesting a significant perturbation in lipid metabolism within ISCs in the context of H3K36me3 loss (Extended Data Fig. 3E). Collectively, these data indicate that H3K36me3 loss perturbs lipid metabolism in ISCs.

Considering that FAO is associated with ISC senescence^15^, we hypothesized that the metabolic disorder of FAO resulting from H3K36me3 deficiency contributes to senescence in ISCs. To provide support evidence for our hypothesis, we analyzed our previously described RNA-seq data and revealed an upregulation of cell senescence markers, including senescence-associated secretory phenotype (SASP) genes and DNA repair factors (Extended Data Fig. 3F). Consistent with these findings, H3K36me3 loss led to a significant increase in the number of SABG-positive cells (Fig. 3G) and the upregulation of senescence markers (Fig. 3H), confirming the role of H3K36me3 in maintaining stem cell function.

### H3K36me3 loss induces a genome-wide alteration in chromatin accessibility

To elucidate whether the increased transcriptional activity of FAO and cell senescence-related genes is a consequence of an altered epigenetic landscape following the loss of H3K36me3, we performed an assay for transposase-accessible chromatin using sequencing (ATAC-seq) on *Setd2*^ISC-KO^ mice. The ATAC-seq analysis revealed genome-wide shifts in chromatin accessibility at approximately 8,683 sites, representing 13.2% of the total reproducible peaks (FDR < 0.05, FC > 1.5) (Supplementary Table 2). Following the depletion of H3K36me3, 64.04% of the affected ATAC-seq peaks exhibited increased accessibility and 35.96% exhibited decreased accessibility (Fig. 4A). The differentially accessible regions were predominantly located at promoters (44.3%), followed by introns (28.6%), intergenic regions (24.7%) and exons (2.9%) (Fig. 4B-C). GO analysis of genes associated with the open chromatin regions in *Setd2*^ISC-KO^ mice also highlighted active biological processes related to lipid metabolism (Extended Data Fig. 4A).

**Figure 4.**
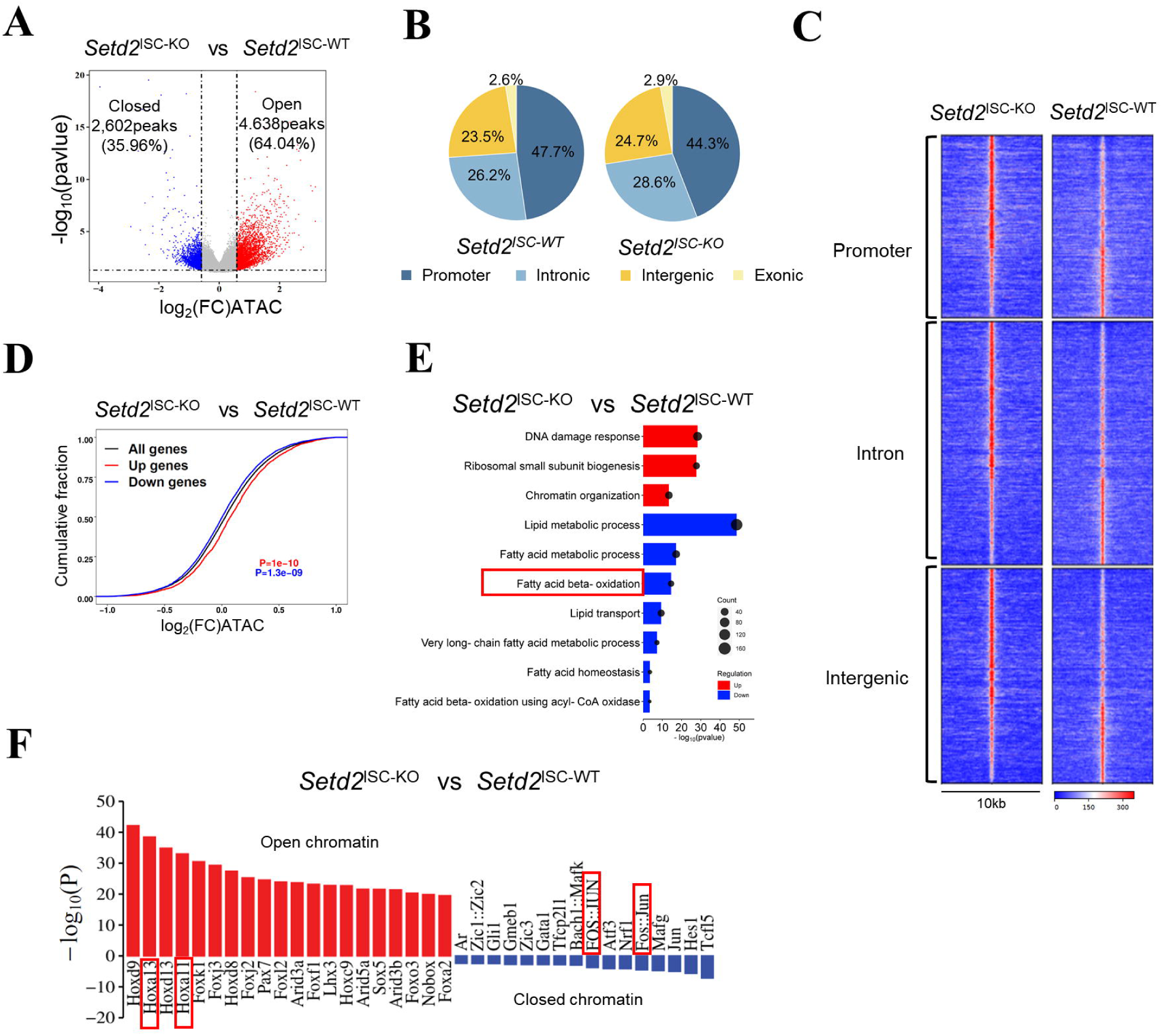
H3K36me3 loss induces a genome-wide alteration in chromatin accessibility. **A.** Volcano plot of ATAC-seq peaks comparing *Setd2*^ISC-KO^ to *Setd2*^ISC-WT^. Peaks with differential chromatin accessibility following SETD2 ablation in ISCs (FDR < 0.05; n = 55,845) are highlighted. The number of peaks with significant changes (FDR < 0.05 and log2 (FC) > 0.585; n = 8,683 peaks) following SETD2 ablation is shown. **B.** Pie charts show the percentage of differentially accessible ATAC-seq peaks (FDR < 0.05) at the promoter, intronic, intergenic, and exonic regions. **C.** Heatmap of differentially accessible ATAC-seq peaks described in an (FDR < 0.05 and log2 (FC) > 0.585) in a 5-kilobase (kb) window grouped by localization at promoter, intron and intergenic regions. **D.** Distribution of chromatin accessibility changes associated with significantly upregulated (red) or downregulated (blue) genes comparing SETD2 ablation with control ISCs. **E.** The GO analysis for the overlapping genes identified in RNA-seq (p < 0.05 and FC > 1.5) and ATAC-seq (FDR < 0.05 and log2 (FC) > 0.585) data are shown, comparing *Setd2*^ISC-KO^ to *Setd2*^ISC-WT^. **F.** Significantly enriched transcription factor binding motifs in open (red) and closed (blue) chromatin peaks comparing *Setd2*^ISC-KO^ to *Setd2*^ISC-WT^. FC, fold change. FDR, false discovery rate.

We subsequently explored the impact of H3K36me3 deficiency on gene expression through chromatin alterations. An overlap was identified between differentially expressed genes and those located near differentially accessible regions in *Setd2*^ISC-KO^ mice (Extended Data Fig. 4B). By linking each open chromatin area to its proximate genes, we established a correlation between changes in chromatin accessibility and gene expression. Upregulated genes were associated with more open chromatin, while downregulated genes were linked to more closed chromatin (Fig. 4D). GO analysis of genes common to both RNA-seq and ATAC-seq data revealed an enrichment of pathways related to lipid metabolism, including fatty acid metabolic processes, lipid storage, fatty acid beta-oxidation, and lipid transport (Fig. 4E). Notably, key genes involved in FAO were found to be downregulated in *Setd2*^ISC-KO^ mice, correlating with changes in chromatin accessibility (Extended Data Fig. 4C).

Open chromatin regions are amenable for transcription factor (TF) binding to initiate a variety of transcription regulation. To identify the transcription factors that regulate senescence-associated gene expression in the absence of H3K36me3, we conducted a motif analysis on the differentially accessible peaks identified by ATAC-seq. The deficiency of H3K36me3 in ISCs resulted in open chromatin regions that were significantly enriched for binding motifs of the Hoxa family of TFs, particularly *Hoxa11* and *Hoxa13* (Fig. 4F). Of note, it has been reported that these specific Hox genes exhibit significant alterations during aging^30^. The closed chromatin regions following H3K36me3 loss were enriched for AP1 (JUN/FOS) transcription factor binding motifs (Fig. 4F), which are known to regulate lipid metabolism-related genes and processes^31^. This may help explain our observations that the absence of H3K36me3 led to the downregulation of FAO. Collectively, our data reveal that loss of H3K36me3 increases chromatin accessibility, enhancing the transcriptional activity of FAO and cell senescence-related genes in ISCs.

### H3K36me3 loss induces epigenetic remodeling and the activation of enhancers

Building on our findings that loss of H3K36 methylation enhances chromatin accessibility and modulates the transcriptional activity, we delved deeper to test whether the distribution of histone post-translational modifications (hPTMs) change. To this end, we employed CUT & Tag to assess both active (H3K27ac, H3K4me1, H3K4me3) and repressive (H3K27me3, H3K9me3) chromatin states in *Setd2*^ISC-KO^ mice. We integrated ATAC-seq with CUT & Tag to examine the genome-wide relationship between chromatin accessibility and active promoter activity, particularly in regions with increased accessibility due to H3K36me3 loss.

Our analysis showed that H3K36me3 deficiency resulted in broad alterations to histone marks, particularly affecting H3K27me3 and H3K4me3, with lesser impacts on H3K4me1 and H3K27ac (Fig. 5A and Supplementary Table 3). H3K36me3 loss significantly altered enhancer marks H3K27ac and H3K4me1 at approximately 700 and 5,000 sites (FDR < 0.05 and log2(FC) > 1) (Fig. 5B), with the majority showing increased signals in H3K36me3-depleted ISCs (Fig. 5B and Extended Data Fig. 5A). This was consistent with our findings of a notable increase in the proportion of cells positive for H3K4me1 and H3K27ac in ISCs deficient in SETD2 (Extended Data Fig. 5B). H3K36me3 deficiency also induced significant changes in H3K4me3 at about 1,300 sites, with 76.97% showing increased signals (Fig. 5B). A positive correlation was noted between changes in ATAC-seq for H3K27ac, H3K4me1 and H3K4me3 (Pearson correlation coefficient of 0.68, 0.62, and 0.70, respectively) (Fig. 5C). Moreover, the loss of H3K36me3 triggered substantial alterations in the repressive H3K27me3 mark at approximately 2,100 sites, with the majority exhibiting increased signals (Fig. 5B). This observation is consistent with the elevated levels of H3K27me3 observed in SETD2-deficient ISCs (Fig. S5B).

**Figure 5.**
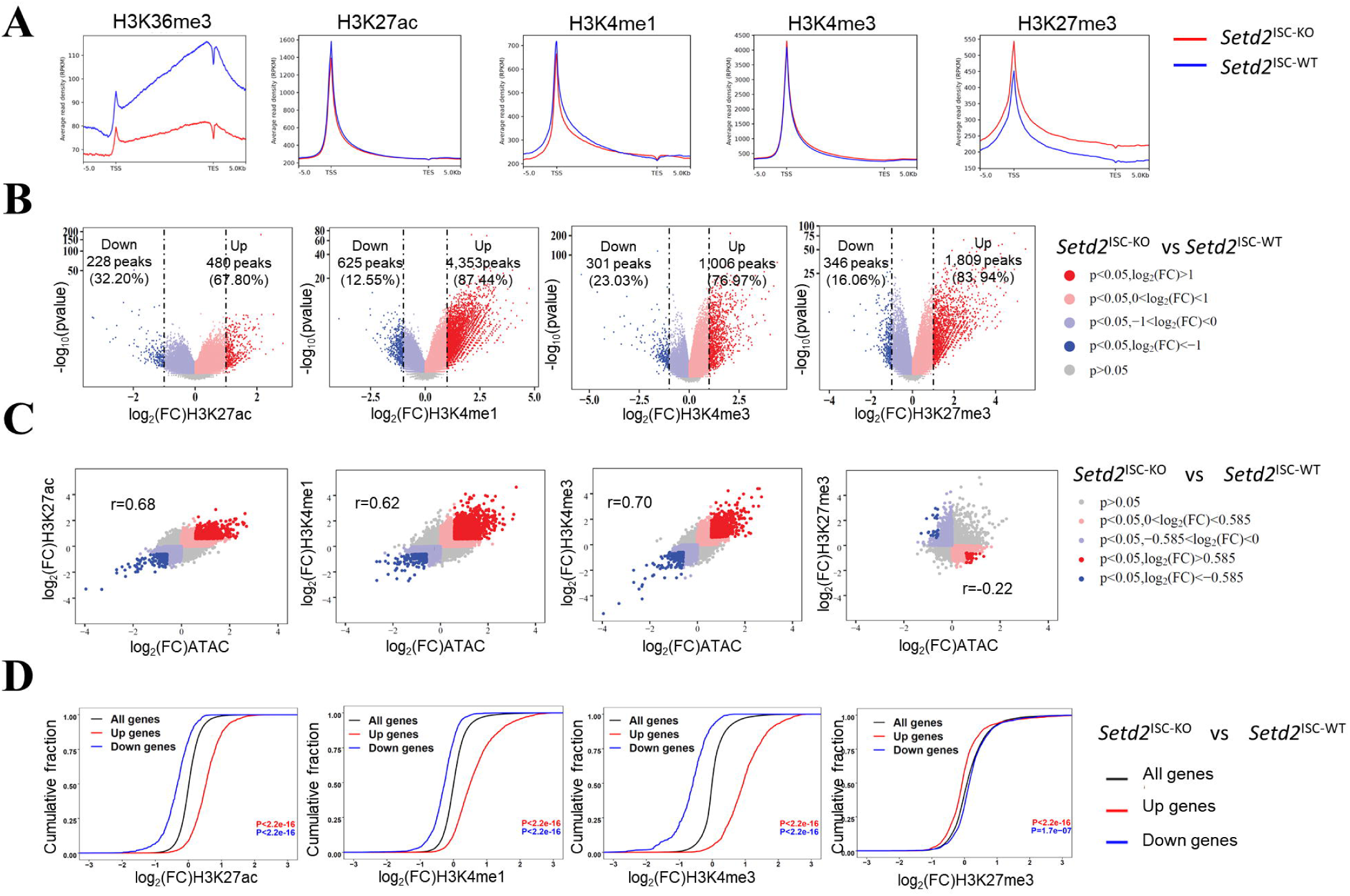
H3K36me3 loss induces epigenetic remodeling and the activation of enhancers. **A.** Metaplots showing the normalized average levels of H3K36me3, H3K27ac, H3K4me1, H3K4me3, and H3K27me3 across gene bodies comparing *Setd2*^ISC-KO^ to *Setd2*^ISC-WT^ ISCs by CUT&Tag; TSS, transcription start site; TES, transcription end site. **B.** Volcano plots showing changes in CUT&Tag for the indicated histone modifications comparing *Setd2*^ISC-KO^ to *Setd2*^ISC-WT^ ISCs. Peaks with differential enrichment for each histone modification (FDR < 0.05) are highlighted. The number of peaks with significant changes (FDR < 0.05 and log2 (FC) > 0.585) in each histone modification is shown. **C.** Scatter plots showing a correlation between log2 (FC) of CUT&Tag for each histone modification and log2 (FC) of ATAC-seq comparing *Setd2*^ISC-KO^ to *Setd2*^ISC-WT^ ISCs. Peaks with significant changes (FDR < 0.05) in histone modification and chromatin accessibility are highlighted. **D.** Cumulative distribution of histone modification changes in significantly upregulated (red) or downregulated (blue) genes (FDR < 0.05) comparing *Setd2*^ISC-KO^ to *Setd2*^ISC-WT^ ISCs. P values were calculated using a one-sided KS test comparing peaks associated with differentially expressed genes to all genes.

Integrated transcriptome and epigenome analyses revealed that upregulated genes gained H3K4me3, H3K4me1, and H3K27ac, while repressed genes showed decreased levels of these marks (Fig. 5D). Notably, CUT&Tag data revealed undetectable H3K9me3 signals (Extended Data Fig. 5A-B), which is consistent with our immunofluorescence staining results that indicate extremely low levels of H3K9me3 in ISCs (Extended Data Fig. 5B). Collectively, our data indicate that the deficiency of H3K36 methylation results in a genome-wide increase in active histone marks associated with enhanced chromatin accessibility.

### H3K36me3 loss activates the chromatin remodeling complex, thereby inducing ISC senescence

Given that chromatin accessibility is primarily modulated by various chromatin-remodeling complexes, we sought to determine whether the loss of H3K36me3 activates these complexes. Our analysis of RNA-seq data from *Setd2*^ISC-WT^ and *Setd2*^ISC-KO^ mice revealed an upregulation of genes associated with chromatin remodeling (Fig. 6A and Extended Data Fig. 6A). This finding is in line with previous research indicating that functional crosstalk between H3K36 methylation and the BRG1/BRM-associated factor (BAF, also called SWI/SNF) chromatin remodeling complex serves as a key regulatory mechanism of transcriptional control during cell fate decisions^32–34^. Intriguingly, the core ATPase subunits, especially SMARCA4, which provide the energy for SWI/SNF-mediated chromatin remodeling, exhibited increased expression levels following the loss of H3K36me3 (Fig. 6B-C). This suggests that the loss of H3K36me3 may activates the SWI/SNF complex, thereby influencing the transcriptional activities in ISCs.

**Figure 6.**
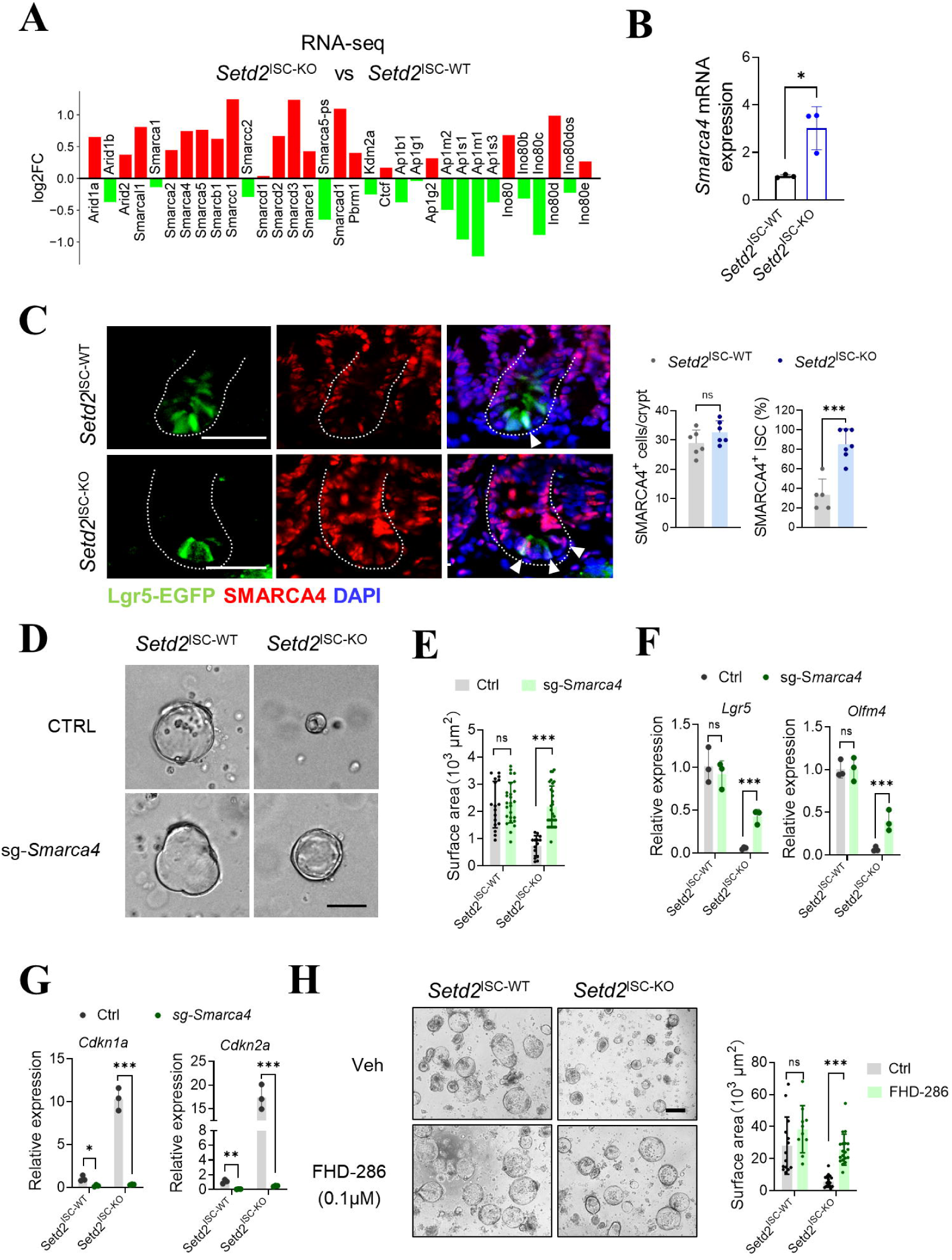
H3K36me3 loss activates the chromatin remodeling complex, thereby inducing ISC senescence. **A.** Normalized bulk RNA-seq counts of genes associated with chromatin remodeling complex comparing *Setd2*^ISC-KO^ to *Setd2*^ISC-WT^ ISCs. **B.** Expression of *Smarca4* in ISCs from *Setd2*^ISC-WT^ and *Setd2*^ISC-KO^. **C.** Representative images and quantification of SMARCA4 in ISCs (EGFP) from *Setd2^ISC-WT^* and *Setd2*^ISC-KO^. **D.** Representative images of organoids formed by crypts of *Setd2^ISC-WT^* and *Setd2*^ISC-KO^, taken 2 days after exposure to either control or sg-*Smarca4* lentivirus. Scale bars, 200 μm. **E.** Quantification was done by measuring the diameter of the spheroids or counting crypt domain formation in **D.**. **F-G.** Expression of stem cell markers (**F**) and cell senescence-related genes (**G**) in organoids in **D.**. **H.** Organoids derived from crypts of *Setd2^ISC-WT^* and *Setd2*^ISC-KO^ mice were treated with DMSO or FHD-286 for 24 hours and then transferred to a basic culture medium. Representative images and quantification are presented on day 3. Scale bars, 200 μm. Quantification was done by measuring the diameter of the spheroids or counting crypt domain formation. For qRT-PCR, GAPDH was used as an internal control. The statistical data represent mean ± s.d. (n = 3 mice per genotype). Student’s t-test: *P < 0.05. **P < 0.01. ***P < 0.001. All images are representative of n = 3 mice per genotype.

Based on these findings, SMARCA4 may be activated following the loss of H3K36me3, thereby recruiting the SWI/SNF complex to enhance chromatin accessibility. We subsequently analyzed the impact of SMARCA4 depletion in organoids derived from *Setd2*^ISC-KO^ mice. Depletion of SMARCA4 in *Setd2*^ISC-KO^ mice restored the organoid-forming capacity of crypts (Fig. 6D and Extended Data Fig. 6B-C) and upregulated the transcription levels of Lgr5 stem cell markers (Fig. 6F). Furthermore, we observed a downregulation of cell senescence markers following SMARCA4 depletion (Fig. 6G).

To further elucidate the role of SMARCA4 in inducing senescence in *Setd2*-deficient ISCs, we employed the SMARCA4 ATPase inhibitor FHD-286^35,36^ (Foghorn Therapeutics), a small molecule that is orally bioavailable and selectively targets BRG1 and BRM ATPase activity. Treatment with FHD-286 increased both the number and size of intestinal organoids derived from *Setd2*^ISC-KO^ mice (Fig. 6H). Consistent with these findings, inhibiting SMARCA4 resulted in increased expression levels of *Lgr5* and *Olfm4*, along with decreased expression levels of cell senescence markers in organoids from *Setd2*^ISC-KO^ mice (Extended Data Fig. 6D-E). Collectively, these data demonstrate that activation of the SWI/SNF complex, induced by H3K36me3 loss, is essential for promoting chromatin remodeling to induce senescence in ISCs.

### Metabolic intervention inhibits the senescence of intestinal stem cells induced by H3K36me3 deficiency

In light of the observation that H3K36me3 loss disturbed FAO metabolism, which is strongly related to cell senescence^37–40^, we were prompted to determine whether metabolic intervention can confer inhibiting effects against H3K36me3 loss-induced ISC senescence. We employed WY14643, a pharmacological agonist of PPARα, to activate the FAO process (Fig. 7A). Crypts from *Setd2*^ISC-KO^ mice treated with WY14643 displayed enhanced outgrowth, budding, and organoid-forming capabilities when cultured ex vivo (Fig. 7B-D). Subsequently, we treated *Setd2*^ISC-KO^ mice with WY14643 and observed that FAO activation significantly rescued the reduction in ISC numbers (Fig. 7E-F) and elevated the expression of ISC markers in *Setd2*^ISC-KO^ mice (Fig. 7G). Furthermore, treatment with WY14643 resulted in a decrease in the number of SABG-positive cells and the expression levels of senescence markers in *Setd2*^ISC-KO^ mice (Fig. 7H-I). These results suggest that activating FAO can significantly impact ISC function and mitigate the ISC senescence process due to H3K36me3 loss.

**Figure 7.**
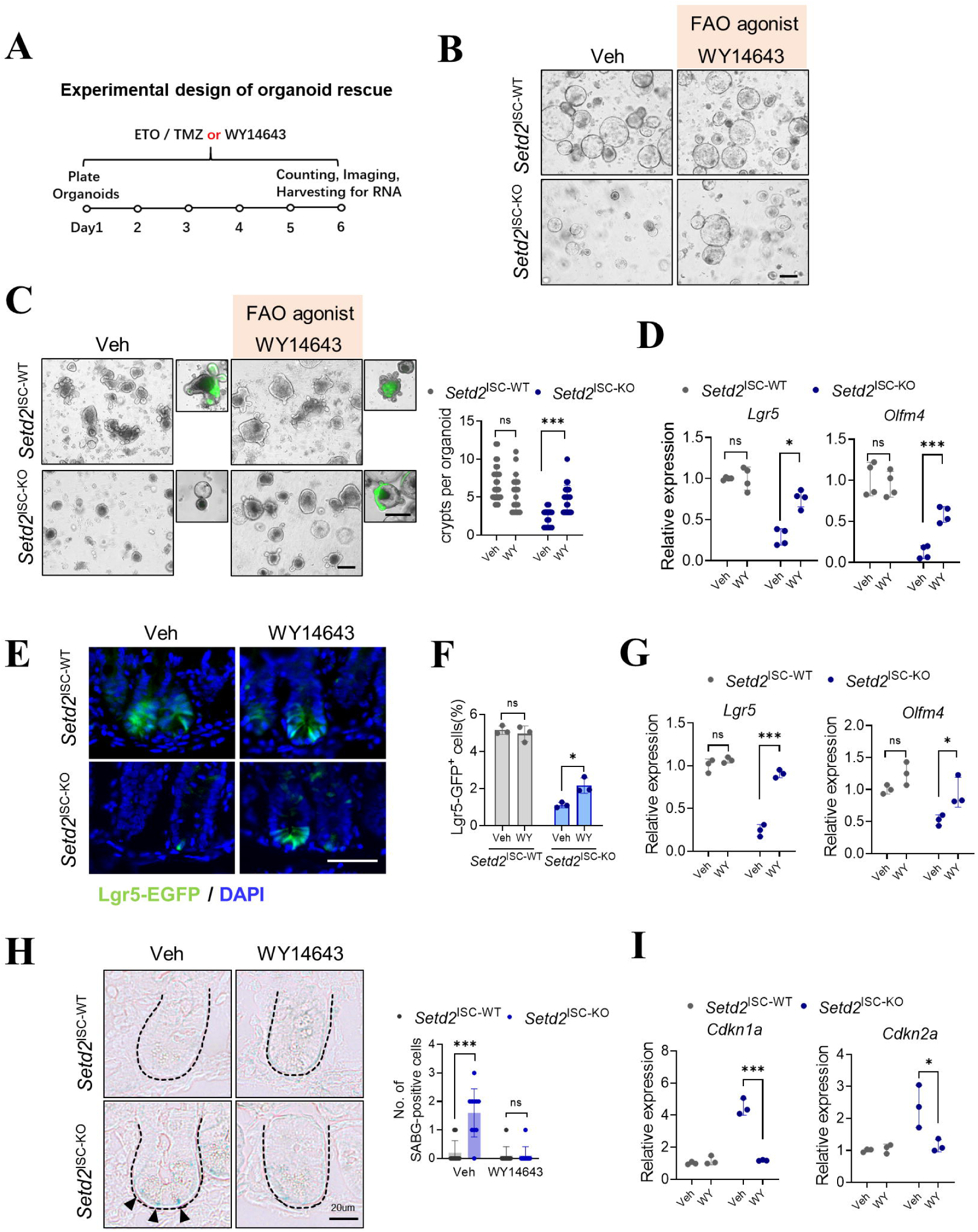
Metabolic intervention inhibits the senescence of intestinal stem cells induced by H3K36me3 deficiency. **A.** A schematic for experimental design of organoid rescue. **B.** Representative morphology (n = 3 independent experiments) of spherical organoids arising from crypts from *Setd2*^ISC-WT^ and *Setd2*^ISC-KO^ mice with or without WY14643 treatment. n = 3 mice per group. Scale bar, 200 μm. **C.** Representative organoid images and quantification of crypts per organoid with or without WY14643 treatment. **D.** qRT-PCR shows a significant increase in the transcript levels of *Lgr5* in the organoids treated with WY-14643. **E.** Representative graphs of crypts containing Lgr5-EGFP+ ISCs in the intestinal sections from *Setd2*^ISC-WT^ and *Setd2*^ISC-KO^ treated with vehicle or WY14643 (n = 3). Scale bars, 50 μm. **F.** Frequency of 7-AAD-Epcam^high^CD24^low^Lgr5^high^ ISCs and within total DAPI-Epcam+ epithelial cells isolated from the small intestine from *Setd2*^ISC-WT^ and *Setd2*^ISC-KO^ treated with vehicle or WY-14643 (n = 3). **G.** Expression of stem cell markers in ISCs from *Setd2*^ISC-WT^ and *Setd2*^ISC-KO^ treated with vehicle or WY14643 (n = 3). **H.** Representative images showed decreased β-SA-Gal positive senescent cells in *Setd2*^ISC-KO^ mice treated with WY14643. n = 3 mice per group. Scale bar, 20 μm. **I.** Expression of cell senescence-related genes in ISCs in **H.** (n = 3). For qRT-PCR, GAPDH was used as an internal control. The statistical data represent mean ± s.d. (n = 3 mice per genotype). Student’s t-test: ns, non-sense. *P < 0.05. **P < 0.01. ***P < 0.001. All images are representative of n = 3 mice per genotype.

## Discussion

Our study elucidates a novel mechanism by which H3K36me3 deficiency promotes ISC senescence. We demonstrated that the loss of H3K36me3 in ISCs creates a permissive epigenetic landscape, which subsequently perturbs lipid metabolism and triggers ISC senescence (Fig. 8). Through an integrative analysis of RNA-seq, ATAC-seq and CUT&Tag data, we uncovered an epigenetic regulatory model in which the depletion of SETD2-caused H3K36me3 results in an altered epigenetic landscape consisting of the activation of the SWI/SNF complex, increased chromatin accessibility and active/permissive histone modifications, thereby facilitating the transcriptional amplification of lipid metabolism and ISC senescence (detailed in Fig. 8). Our findings underscore the significance of H3K36me3 and FAO in ISC senescence and suggest that targeting either of them may provide a strategy to maintain intestinal homeostasis and develop treatments for age-related intestinal diseases and stem cell dysfunction.

**Figure 8.**
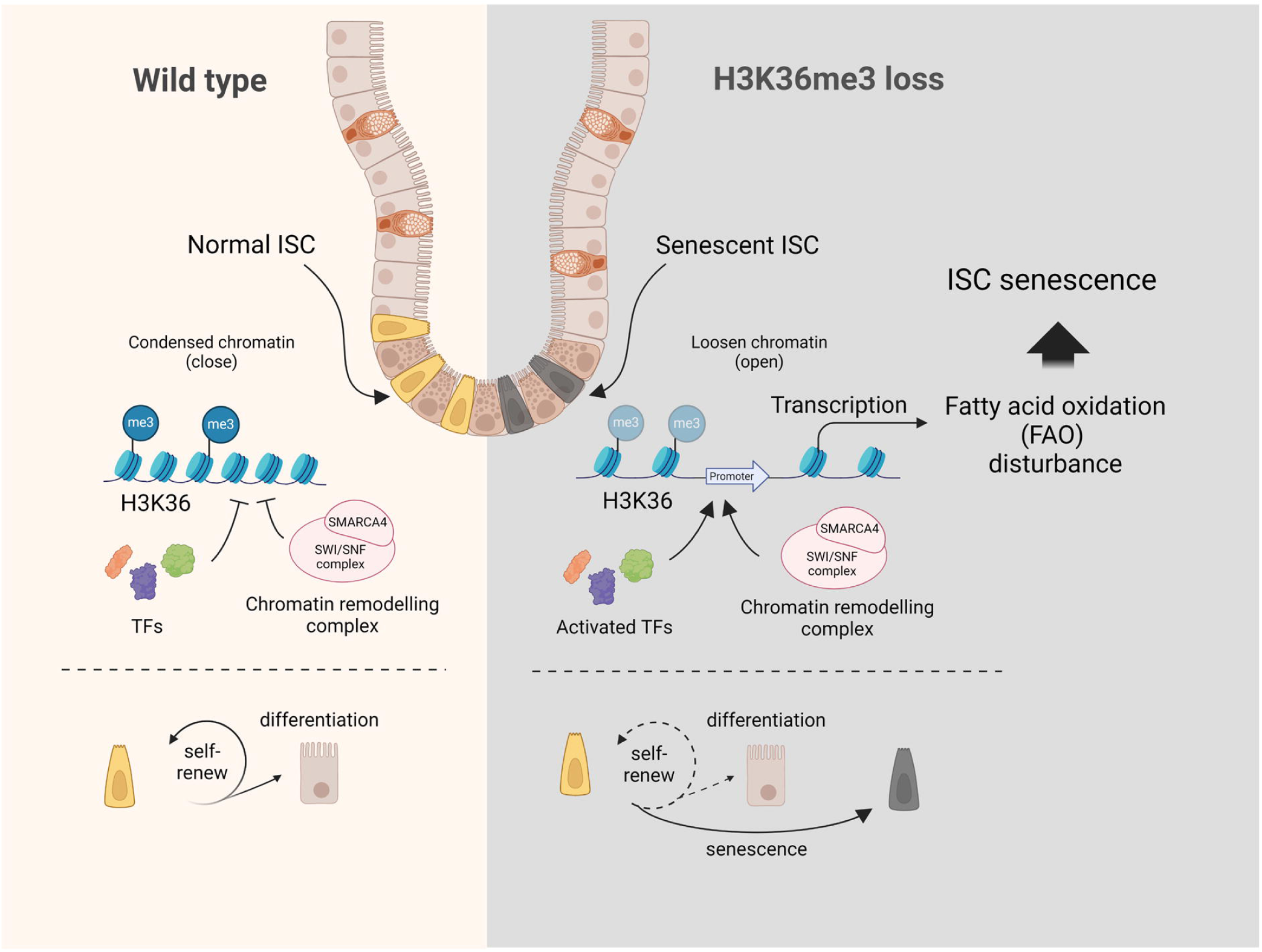
**A schematic summarizing the model of H3K36me3 deficiency promotes ISC senescence via disturbing lipid metabolism**

The implications of epigenetic dysregulation vary by tissue type and specific epigenetic factors, leading to a spectrum of outcomes that can range from subtle alterations in the differentiation potential of affected stem cells to severe consequences such as stem cell depletion. While limited histone modifications have been shown to uniquely contribute to the preservation of ISC identity ^19,21,22,41^, our current study further demonstrates that H3K36me3 is a key determinant in maintaining ISC stemness. This provides novel insights, building on our previous study where we identified H3K36me3 as a central player in maintaining intestinal epithelial homeostasis^42–45^. Here, we reveal that H3K36me3 deficiency impairs ISC proliferation and differentiation and results in ISC senescence characterized by upregulated cell senescence markers and increased SA-β -Gal levels. Mechanistically, the loss of H3K36me3 in ISCs creates a permissive epigenetic landscape, which facilities the transcriptional activity of genes associated with cellular senescence, including cell cycle related gene, SASP genes, cell surface markers and DNA repair factors. Our current study provides new insights into the role of H3K36me3 in the senescence of ISC.

An intriguing finding in the current study is the identification of FAO metabolism as a mediator between histone modification and ISC senescence. Specifically, the deficiency of H3K36me3 inhibits FAO metabolism. This discovery is particularly significant, as our data demonstrate that disturbances in FAO metabolism can impair the biosynthetic and bioenergetic processes essential for ISC proliferation and survival, ultimately contributing to ISC senescence. Of note, in the present work, the senescence of ISCs caused by H3K36me3 loss was alleviated through metabolic intervention. In support of our findings, previous research has highlighted the role of FAO metabolism in the long-term ISC maintenance^46–48^. Therefore, our study underscores the importance of understanding and preserving FAO metabolism for the health and function of ISCs, particularly in the context of aging.

It’s worth emphasizing that in the present study, we delineated a novel epigenetic mechanism through which the loss of H3K36me3 leads to an upregulation of SMARCA4, a central component of the SWI/SNF complex, which further induces chromatin remodeling and disturbs the epigenetic landscape. This is in line with previous research indicating that the functional crosstalk between and the SWI/SNF complex serves as a key regulatory mechanism for transcriptional control during cell fate decisions^32–34^. Interestingly, the depletion or inhibition of SMARCA4 significantly restores the self-renewal capacity of ISCs in the context of H3K36me3 loss, suggesting a complex co-regulation among epigenetic factors. Thus, our findings underscore the significance of histone modification marks in activating chromatin remodeling factors, such as the SWI/SNF complex.

Collectively, our data demonstrate that the loss of H3K36me3 induces an altered epigenetic state in ISCs through the regulation of chromatin-remodeling complexes, specifically impacting lipid metabolism and promoting cell senescence. This study highlights the intricate interplay between histone modifications and lipid metabolism within the context of stem cell biology, providing insights into the epigenetic foundations of intestinal homeostasis and disease.

## Methods

### Experimental models and study participant details Mice

All mice were maintained in a specific-pathogen-free (SPF) facility, and all experimental protocols involving mice were approved by the Renji Hospital Animal Care and Use Committee (202201027). SETD2-flox mice (*Setd2*^f/f^) were generated as previously reported^49,50^. Lgr5-EGFP-CreERT2 mice were gifted by Prof. Shan Sun at Fudan Univeristy in Shanghai. These strains were interbred to generate the experimental cohorts, which included the following genotypes: Lgr5-EGFP-CreERT2; *Setd2*^f/f^ (*Setd2*^ISC-WT^ and *Setd2*^ISC-KO^). Mice were harvested at specified time points for intestinal histological investigation. Mice that lost more than 20% of their body weight within one week were euthanized and recorded as deceased. All mice were maintained on a C57BL/6J background, and littermates receiving the same treatment were used as controls in the experiments.

### Pharmacological studies in mice Intraperitoneal injection

Palmitic acid (also called WY-14643; MCE, HY-16995) was administered to mice at a dose of 1 mg/kg/day by intraperitoneal injection (i.p.), using 0.1% DMSO in normal saline as the vehicle control^51^. Etomoxir (MCE, HY-50202) was administered to mice at a dose of 15 mg/kg/day by i.p. injection, also using 0.1% DMSO in normal saline as the vehicle control^52^. Trimetazidine (MCE, HY-B0968) was administered to mice at a dose of 10 mg/kg/day by intragastric gavage (i.g.), using PBS as the vehicle control.

### In vivo tamoxifen treatment

To delete *Setd2* in lgr5^+^ISCs, adult Lgr5-EGFP-CreERT2; *Setd2*^f/f^ mice were intraperitoneally injected with tamoxifen (SIGMA, T5648) suspended in corn oil at a concentration of 10 mg/ml. The dosage administered was 5 μl per gram of body weight, given daily for five consecutive days, as previously described^48^. Control mice received intraperitoneal injections of corn oil.

### Detection of fluorescent cells in tissues

To detect Lgr5-EGFP+ cells, we fixed mouse intestinal tissues in 4% paraformaldehyde (PFA) in phosphate-buffered saline (PBS) at 4°C for 1h and subsequently rotated them overnight at 4°C in 30% sucrose in PBS. The tissues were then embedded in OCT compound (Tissue-Tek, 4583) and stored at −80°C. Tissue sections (10 μm) adhered to glass slides were washed in PBS and mounted in 1% BSA-PBS containing 4′,6-diamidino-2-phenylindole (DAPI; Vector Laboratories, H-1200). Fluorescent cells were visualized and counted, and images were analyzed by ImageJ software.

### Immunofluorescence

Intestinal tissue sections were incubated overnight with the following antibodies: H3K36me1 (Abcam, ab9048, 1:1000), H3K36me3 (Abcam, ab9050, 1:1000), H3K27ac (Abcam, ab4729, 1:1000), H3K27me3 (Abcam, ab6002, 1:1000), H3K4me1 (Abcam, ab8895, 1:1000), H3K4me3 (Abcam, ab8580, 1:1000), H3K9me3 (Abcam, ab8898, 1:1000), and SMARCA4 (Abcam, ab110641, 1:1000) at 4°C. This was followed by incubation with either Alexa Fluor 594-conjugated anti-rabbit (Jackson ImmunoResearch, 111-585-003, 1:500) or Alexa Fluor 594-conjugated anti-mouse (Jackson ImmunoResearch, 115-585-003, 1:500) antibodies for 1 h at room temperature. Subsequently, the sections were stained for 10 minutes with 4′,6-diamidino-2-phenylindole (DAPI; 1 μg/ml). Images were captured using a microscope.

### LipidTox staining

Mouse intestines were cut and prefixed in fresh 4% paraformaldehyde (PFA) for at least two hours at room temperature, then soaked and washed three times in a 30% sucrose-PBS solution. Prefixed tissue was snap-frozen in an OCT compound (Sakura) and 10 μm sections were cut and mounted on glass slides. Slides were fixed in 4% fresh PFA for 20 min and washed twice with PBS. Fixed slides were stained for 2 h in the dark with LipidTox (Thermo Fisher H34476) working solution (1:500 diluted in PBS), followed by staining for 10 min with 4′,6-diamidino-2-phenylindole (DAPI; 1 μg/ml). Images were captured using a microscope.

### Lipid quantification in intestinal epithelial cells

A total of 5-10 × 10^6^ cells were isolated from mouse intestinal crypts and subjected to lipid extraction using a Lipid Extraction Kit (Cell Biolabs STA-162). The extracted lipids were air-dried, resuspended in 200 µl of cyclohexane, and quantified using a Lipid Quantification Kit (Cell Biolabs STA-613).

### SA-β-Gal staining

SA-β-Gal staining was conducted using the Senescence β-Galactosidase Staining Kit (Cell Signaling Technology, 9860S). The procedure followed the manufacturer’s instructions provided in the kit. Briefly, intestinal sections were fixed in 1X Fixative Solution for 15 minutes and then washed twice with phosphate-buffered saline (PBS). The fixed slides were incubated with the β-Galactosidase Staining Solution overnight. While the β-galactosidase was still present on the slides, positive cells were examined under a microscope to observe the development of blue color. For long-term storage, the β-Galactosidase staining solution should be removed, and the slides should be overlaid with 70% glycerol. Store the slides at 4°C.

### Intestinal crypt isolation and crypts-derived organoids culture

The culture of organoids derived from intestinal crypts was adapted from previous reports^53^. The basic culture medium consisted of advanced DMEM/F12 (Gibco, PHG0312L) supplemented with penicillin/streptomycin (InvivoGen, ant-pm-1), 10 mM HEPES (Invitrogen, 15630080), 1× B27 (Invitrogen, 17504044), 1× N2 (Invitrogen, A1370701), L-Glutamine (Solarbio, G0200) and N-acetylcysteine (sigma, A9165). Additionally, the medium contained 250 ng/ml R-spondin1, 50ng/ml m-EGF, and 100 ng/ml m-noggin. Small intestines were collected and flushed with ice-cold PBS, then incubated in 10mM EDTA at 4 °C for 20 minutes. Tissues were subsequently washed with PBS. Crypts were mechanically separated from tissues by pipetting up and down, followed by filtration through a 70-μm cell strainer. This mechanical separation was repeated 3 to 5 times until purified crypts were isolated and counted. The isolated crypts were then embedded in a 1:1 mixture of Matrigel (Corning 356231, growth factor reduced) and culture medium at a concentration of 5 to 10 crypts per μl. Intestinal crypts were plated in 10 to 50 μl droplets onto the flat bottom of Corning 48- and 24-well plates and allowed to solidify for 15 minutes in a 37°C incubator. Subsequently, 150 to 500 μl of crypt culture medium were overlaid onto the Matrigel in each well, depending on the specific culture plate used. Organoids were exposed to palmitic acid (MCE, HY-16995), trimetazidine (MCE, HY-B0968), etomoxir (MCE, HY-50202), or BRM/BRG1 ATP Inhibitor-1 (MCE, HY-119374). The crypt culture medium was changed every two days and maintained at 37°C in fully humidified chambers containing 5% CO_2_. For serial organoid culture, primary organoids were removed from Matrigel, mechanically dissociated into single-crypt domains, and then transferred to fresh Matrigel. Passaging was performed every 5 to 7 days with a 1:5 split ratio.

### Intestinal crypts single cell isolation and ISCs sorting

In summary, the small intestines were collected and rinsed with ice-cold PBS. They were then cut longitudinally into approximately 2-mm pieces and incubated in 10 mM EDTA at 4°C for 20 minutes. Following this, the pieces were shaken in ice-cold PBS for 1 minute, and this mechanical separation was repeated three times. The resulting mixture was filtered through a 70-μm cell strainer to obtain enriched crypts. The isolated crypts were subsequently incubated in TrypLE Express supplemented with DNase I (800 U/ml) at 37°C for 10 minutes, after which they were filtered through a 40-μm mesh cell strainer to yield single crypt cells. The single crypt cells isolated from Lgr5-EGFP mice were stained with 7-AAD-PercP, CD24-PE-Cy7-561, and Epcam-APC antibodies. Lg5^+^ISCs were sorted as 7AAD^+^Epcam^high^CD24^low^EGFP^high^ with BD FACS Aria II.

### Single intestinal stem cell-derived organoids culture

1*10^4^ cells of Lgr5^+^ISCs were plated per well in 10 μl droplets of Matrigel (Corning 356231) mixed with ISC culture medium in a 1:1 ratio onto the flat bottom of a Corning 24-well cell culture plate. The droplets were allowed to solidify for 10 to 15 minutes in a 37°C incubator. Following this, 150 μl of ISC culture medium was overlaid onto the Matrigel in each well. The ISC culture medium was changed every two days and maintained at 37°C in fully humidified chambers containing 5% CO2. The number of organoids and organoid-forming crypts was counted per well 4 to 9 days after the initiation of cultures. The ISC culture medium was prepared using a basic culture medium supplemented with 1 μM JAGGED-1 (Genscript, RP20331), 10 μM Y-27632 (MCE, HY-10583), 2.5 μM CHIR99021 (Stemgent, 04-0004), and 100 ng/ml Wnt-3a (Peprotech).

### Lentivirus production and the indicated lentivirus-reconstituted organoid formation

Control and sg-*Smarca4* lentiviruses were produced by Hunan Fenghui Biotechnology Co., Ltd. Purified crypts were infected with the specified recombinant lentiviruses at a multiplicity of infection (MOI) of 15. The complexes were then inoculated at 400 × g and 20 °C for 1 h, plated hour, followed by plating in a Corning 24-well cell culture plate supplemented with the same volume of Matrigel Matrix (Corning, 356231) and 8 μg/mL polybrene (Santa Cruz Biotechnology, sc-134220), and incubated in a culture incubator. The plates were maintained at 37 °C in 5% CO_2_ with a medium change every 1 days and organoids were analyzed under a light microscope.

### In vitro EdU incorporation assay

To assess epithelial proliferation, mice were injected intraperitoneally (i.p.) with 80 μl of 10 mM EdU (RIBOBIO, C10310) and analyzed 2 hours after injection.

### RNA isolation and RT-qPCR

Total RNA was isolated from sorted-ISCs, dissociated-organoids, or small intestine epithelial cells using TRIzol and subsequently reverse transcribed into cDNA using the High-Capacity cDNA Reverse Transcription Kit. Quantitative reverse transcription polymerase chain reaction (RT-qPCR) analysis was performed using SYBR Green Master Mix and specific primers. The signals were normalized to GAPDH levels within each sample, and the normalized data were used to calculate relative gene expression levels using the ΔΔCt method. Data analysis was performed with three biological replicates. All primer sequences are provided in Supplementary Table 4.

### Organoid measurement and image

Organoid measurements were performed as previously reported^54,55^. The number of organoids and crypts was counted manually, and crypt formation was evaluated by calculating the ratio of crypts to organoids in each well. Organoids were imaged using either an Olympus IX50 Inverted Microscope or an Invitrogen EVOS M500 Inverted Microscope. Unless otherwise specified in the figure legends or method details, organoid assays included 3 to 6 wells per group, with samples analyzed from at least 3 different mice.

### Metabolomics and lipidomics profiling

The sample stored in a -80 °C freezer was thawed on ice. The thawed sample was homogenized by a grinder at 30 Hz for 20 s. A 400 μL solution composed of methanol and water in a 7:3 volume ratio, containing the internal standard, was added to the 20 mg of ground sample, and shaken at 1500 rpm for 5 minutes. After being placed on ice for 15 min, the sample was centrifuged at 12,000 rpm for 10 min (4 °C). A 300 μL aliquot of the supernatant was collected and stored at -20 °C for 30 minutes. The sample was then centrifuged again at 12,000 rpm for 3 min (4 °C). A 200 μL aliquot of the supernatant was transferred for LC-MS analysis. Unsupervised principal component analysis (PCA) was performed using the ‘prcomp’ function in R (www.r-project.org). The data were scaled to unit variance prior to PCA. For the two-group analysis, differential metabolites were identified based on variable importance in projection (VIP) values (VIP > 1) and P-values (P-value < 0.05, Student’s t-test). VIP values were extracted from the OPLS-DA results, which also included score plots and permutation plots generated using the R package MetaboAnalystR. The data were log-transformed (log2) and mean-centered before OPLS-DA. To avoid overfitting, a permutation test with 200 permutations was conducted.

### ATAC sequencing

In brief, we FACS sorted 1*10^5^ Lgr5^+^ISCs each biology samples. Cell samples were sent to Genefund Biotech (Shanghai, China) for ATAC-Seq library preparation and data analysis.

### RNA sequencing

RNA was extracted using Trizol reagent from Life Technologies Corp., and DNase treatment was applied to remove any potential genomic DNA contamination. PolyA mRNA was enriched using the NEBNext PolyA mRNA Magnetic Isolation Module from New England Biolabs, Ipswich, MA, USA, which was then used to prepare RNA-Seq libraries with the NEBNext Ultra Directional RNA Library Prep Kit for Illumina. These libraries were sequenced on an Illumina instrument with a paired-end 150 bp read length protocol. To enhance the accuracy of the sequencing data, we employed Cutadapt (version 1.9.1) and Trimmomatic (version 0.35) to process the raw reads, removing any sequencing adapters, nucleotide sequences shorter than 35 bases, and low-quality reads. The quality of the resulting reads was validated using FastQC. The high-quality reads were then mapped to the mouse reference genome (GRCm38 assembly) with HISAT2 software. Gene expression levels were estimated using the FPKM method with StringTie^56^ , and differential gene expression was analyzed using the Ballgown R package^57^. To correct for multiple testing, we utilized the false discovery rate (FDR) approach to calculate adjusted P-values, which facilitated the determination of the statistical significance of the observed gene expression differences. Genes with an adjusted P-value below 0.05 were considered for further investigation.

### CUT&TAG sequencing

We conducted CUT&Tag analysis utilizing the Hyperactive® Universal CUT&Tag Assay Kit for Illumina (Vazyme, TD903), adhering to the manufacturer’s protocol with minor adjustments. To begin, 100,000 freshly isolated ISCswere used to isolate nuclei, which were then combined with ConA beads and incubated at room temperature for 10 minutes. Subsequently, these complexes were incubated with antibodies targeting H3K36me3, H3K27ac, H3K27me3, H3K4me1, H3K4me3, or IgG as a negative control, followed by the addition of a secondary antibody the following day. A diluted pA-Tn5 adapter complex was introduced to the mixture and incubated for 1 hour at room temperature. After thorough washing, the system was disrupted by the addition of 5 × TTBL. Proteinase K and DNA extraction beads were employed to acquire the purified DNA. The sequencing library was constructed following 12 cycles of PCR amplification and then sequenced on the Illumina Novaseq 6000 platform using a paired-end 2×150 sequencing strategy, as per Genefund Biotech’s (Shanghai, China) guidelines. To ensure high-quality data, raw reads were processed with Trimmomatic v0.38^58^ to remove sequencing adapters, short reads (under 36 bp), and low-quality sequences (non-default parameters: SLIDINGWINDOW:4:15 LEADING:10 TRAILING:10 MINLEN:36). FastQC^59^ was applied with default settings to verify the quality of the reads. The cleaned reads were aligned to the mouse genome (GRCm38 assembly) using Bowtie2 v2.3.4.1^60^ with specific parameters (-I 10 -X 700 --no-discordant --no-mixed --local --very-sensitive-local). Duplicate reads were eliminated using picard MarkDuplicates. MACS v2.1.2^61^ was employed for peak detection with a q-value threshold set at 0.05 (non-default parameters: -f BAMPE -g hs/mm -q 0.05). The ChIPseeker R package^62^ was utilized for the annotation of peak sites relative to gene features. The entire analysis was conducted by Genefund Biotech (Shanghai, China).

## Quantifaction and statistical analysis

### Statistical analysis of biological data

In general, the data are presented as the mean ± SD (standard deviation) from at least three independent repeats. We replicate unpaired employed Student’s tests (t t-tests the assess level and level, considering a as the statistically significant difference. indicative of a samples was excluded from any analysis. Additionally, we presented the Furthermore, morphology provided representative images of from at immunostaining replicates.

## Data availability

All data associated with this study are present in the paper or the Supplementary Materials. RNA-seq, ATAC-seq, and CUT&Tag analysis data have been submitted to GEO under accession numbers: GSE292892, GSE292016, GSE292017 and GSE292018.

## Acknowledgement

This work was supported by funds from National Key R&D Program of China (2022YFA1302704 to L.L. and W-Q.G., 2023YFC1404101 to W-Q.G.), National Natural Science Foundation of China (U23A20441, W2431055 to W.-Q.G., 82372604 to L.L.), the Peak Disciplines (Type IV) of Institutions of Higher Learning in Shanghai, 111 Project (B21024) and KC Wong foundation to W-Q.G. and Interdisciplinary Program of Shanghai Jiao Tong University (YG2024ZD11). We would like to thank Genefund Biotech (Shanghai, China) for their assistance with the data analysis.

## Author contributions

L.L., W.-Q.G. conceived the experimental concept, designed the experiments, and interpreted the data; Y.X. performed most of the experiments and wrote the manuscript; Z.W. helped with sample collection and CUT & Tag experiments; W.F., H.R., D.W., W.Z., R.A. and N.L. assisted in some experiments; W.-Q.G. assisted in some discussion; L.L., provided the overall guidance. All authors read and approved the final manuscript.

## Competing Interests

The authors declare no competing interests.

## Research animals

All experimental procedures were approved by the Institutional Animal Care and Use Committee of Shanghai Jiao Tong University.

## Notes

### Competing Interest Statement

The authors have declared no competing interest.

