## Extended data figure for "Histone 3 lysine 36 trimethylation by SETD2 shapes an epigenetic landscape in intestinal stem cells to orchestrate lipid metabolism and prevent cell senescence"

Extended Data Figure 1.

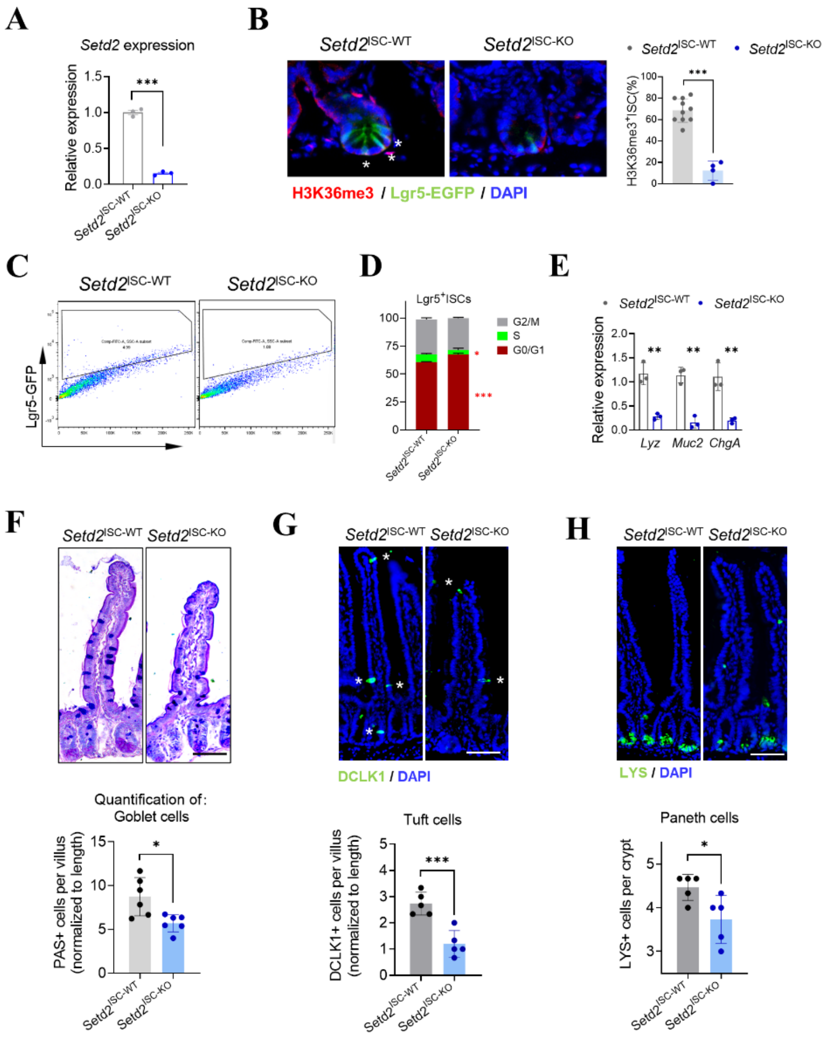


**Extended Data Figure 1. A.** qPCR analysis of *Setd2* expression in ISCs following five consecutive days of tamoxifen injections in *Lgr5-EGFP-CreERT2; Setd2*$^{f/f}$ mice. **B.** Representative images and quantification of H3K36me3 levels (red) in ISCs (green) from *Setd2*$^{ISC\text{-}WT}$ and *Setd2*$^{ISC\text{-}KO}$ mice. **C.** Flow cytometry graphs show the frequency (%) of Lgr5-EGFP+ ISCs in *Setd2*$^{\text{ISC-WT}}$, and *Setd2*$^{\text{ISC-KO}}$ mice after 5 consecutive days of TAM treatment. Data are presented for one pair from n = 3 mice of each genotype. **D.** Cell cycle analysis of Lgr5+ ISCs from *Setd2*$^{\text{ISC-WT}}$, and *Setd2*$^{\text{ISC-KO}}$ mice following five consecutive days of tamoxifen treatment. Data are presented as mean ± SD (n = 3). **E.** qPCR analysis of *Lyz*, *Muc2* and *ChgA* expression in ISCs from *Setd2*$^{\text{ISC-WT}}$, and *Setd2*$^{\text{ISC-KO}}$ mice (n = 3). **F-H.** Representative images and quantification of goblet cells (PAS+), tuft cells (DCLK+), and Paneth cells (LYS+) from *Setd2*$^{\text{ISC-WT}}$ and *Setd2*$^{\text{ISC-KO}}$ mice. Scale bar, 50 µm, n = 3 mice per group. Data are shown as mean ± s.e.m. of biologically independent samples. Statistical significance was determined by a two-tailed Student's t-test. *P < 0.05, **P < 0.01, ***P < 0.001. For qRT-PCR, GAPDH was used as an internal control. All images are representative of n = 3 mice per genotype.

Extended Data Figure 2.

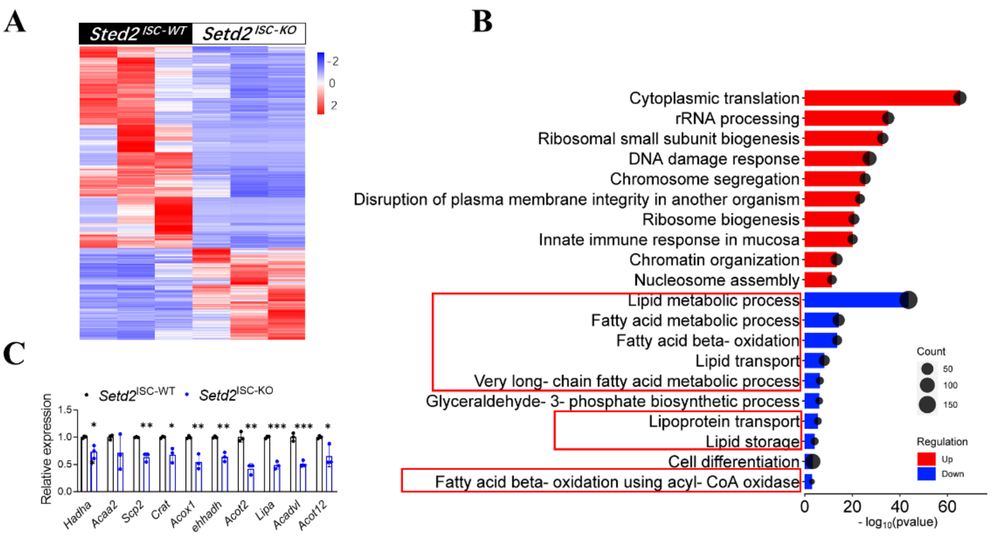


**Extended Data Figure 2. A.** Differential gene expression analysis by RNA-seq in ISCs isolated from *Setd2*$^{\text{ISC-WT}}$ and *Setd2*$^{\text{ISC-KO}}$ mice (n = 3). **B.** Total enriched pathways (GO) in ISCs isolated from *Setd2*$^{\text{ISC-WT}}$ and *Setd2*$^{\text{ISC-KO}}$ mice (n = 3). **C.** qPCR analysis of genes related to β-oxidation in ISCs from *Setd2*$^{\text{ISC-WT}}$ and *Setd2*$^{\text{ISC-KO}}$ mice.

Extended Data Figure 3.

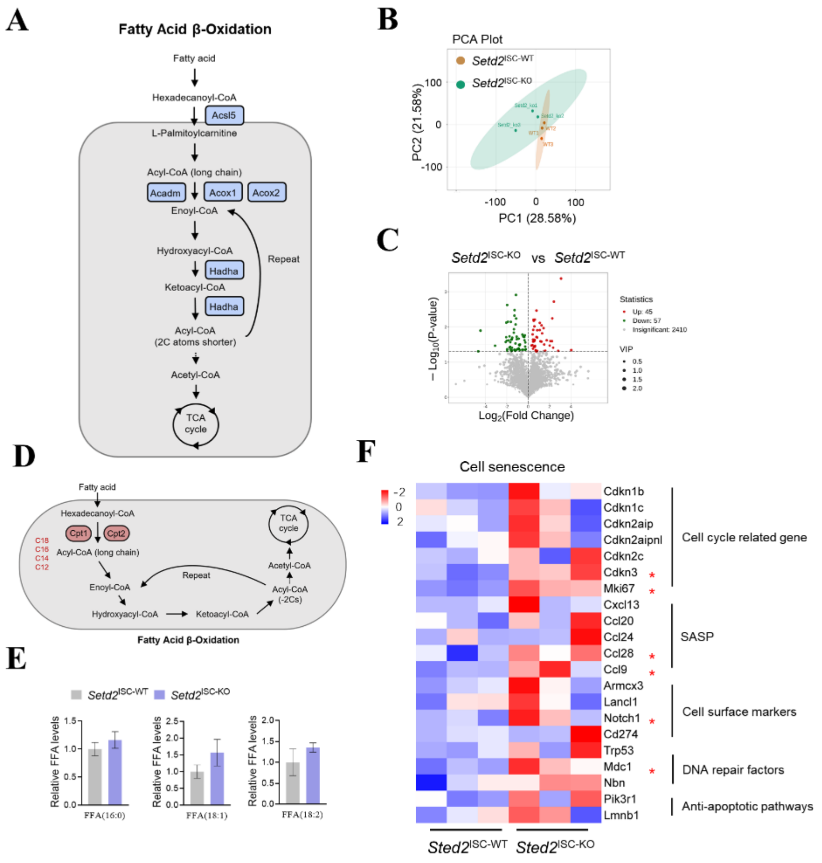


**Extended Data Figure 3. A.** Schema of fatty acid β-oxidation, with the most critical representative upregulated and downregulated genes in *Setd2*$^{\text{ISC-KO}}$ mice highlighted in blue (p-value < 0.05 and fold change >1.5). **B.** Principal-component analysis (PCA) of crypts from *Setd2*$^{\text{ISC-WT}}$ and *Setd2*$^{\text{ISC-KO}}$ mice. **C.** Volcano plot of differential normalized relative abundance of metabolites from *Setd2*$^{\text{ISC-KO}}$ versus *Setd2*$^{\text{ISC-WT}}$ (n = 3). **D.** Schematic illustration of mitochondria β-oxidation system. **E.** Bar charts showing representative FFA levels after SETD2 deletion in ISCs. **F.** Heatmaps of RNA-seq data show increased transcript levels of cell senescence-related genes in *Setd2*$^{\text{ISC-KO}}$ mice (n = 3). Asterisks indicate genes that are significantly increased in *Setd2*$^{\text{ISC-KO}}$ mice. Data are shown as mean ± s.e.m. of biologically independent samples. Statistical significance was determined by a two-tailed Student's t-test. *$P < 0.05$, **$P < 0.01$, ***$P < 0.001$. For qRT-PCR, GAPDH was used as an internal control.

Extended Data Figure 4.

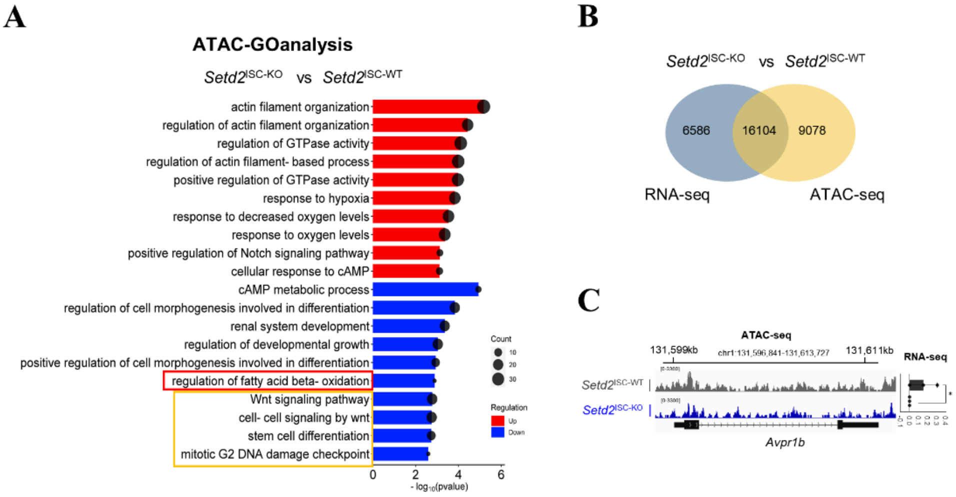


**Extended Data Figure 4. A.** The GO analysis of ATAC-seq data shows the altered pathways following the ablation of SETD2 in ISCs. **B.** Venn diagram showing the overlap between differentially expressed genes and those located near differentially accessible regions in *Setd2*$^{\text{ISC-KO}}$ mice. **C.** Left, IGV tracks of open chromatin at *Avpr1b* (with y-axis scales as indicated). Right, normalized bulk RNA-seq counts of *Avpr1b* in ISCs (n = 4).

Extended Data Figure 5.

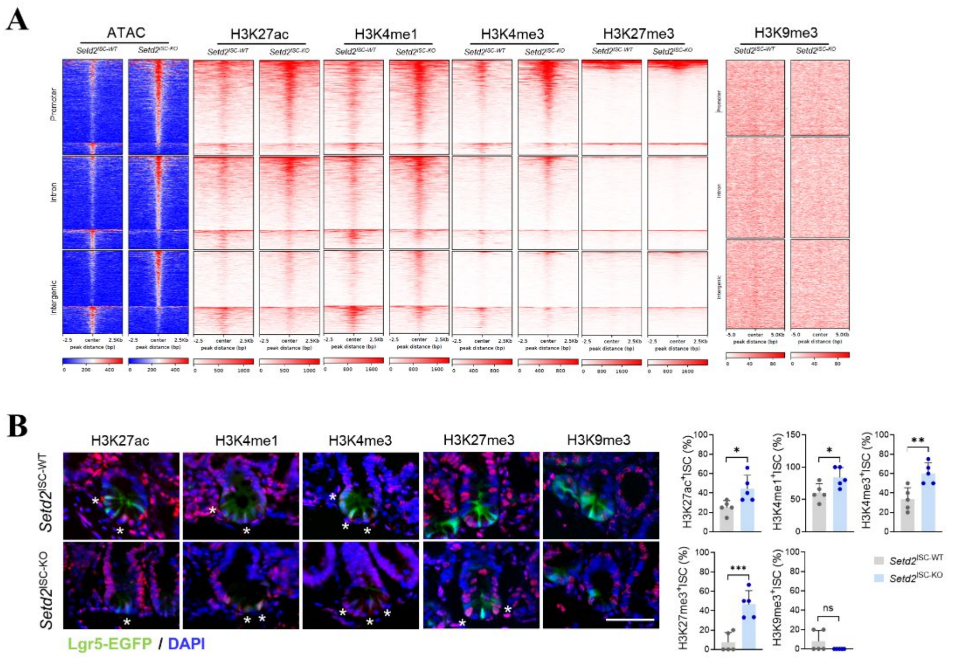


**Extended Data Figure 5. A.** Heat maps of differentially accessible ATAC-seq peaks (FDR < 0.05 and log2 (FC) > 0.585; n = 3,816) in a 5-kb window grouped by localization at the promoter, intron and intergenic regions and CUT&Tag signals for the indicated histone modifications in the same regions of ATAC-seq peaks. **B.** Representative images and quantification of indicated histone modifications (red) in ISCs (green) from *Setd2*$^{ISC\text{-}WT}$ and *Setd2*$^{ISC\text{-}KO}$ mice (n = 3). Data are shown as mean ± s.e.m. of biologically independent samples. Statistical significance was determined by a two-tailed Student's t-test. *P < 0.05, **P < 0.01, ***P < 0.001.

Extended Data Figure 6.

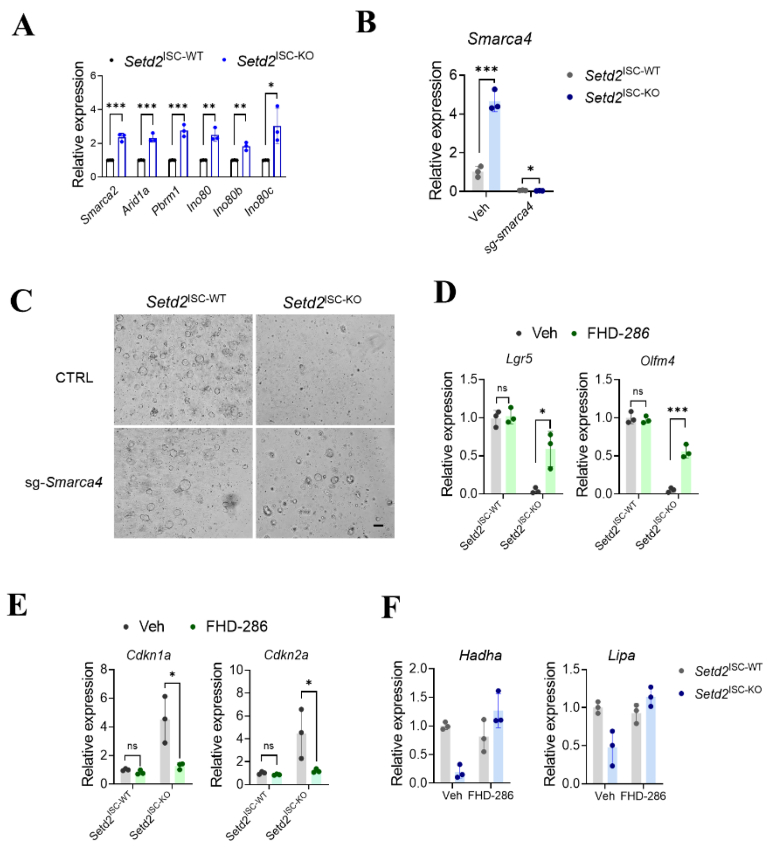


**Extended Data Figure 6. A.** qPCR analysis of genes related to chromatin remodeling complex in ISCs from *Setd2*$^{\text{ISC-WT}}$ and *Setd2*$^{\text{ISC-KO}}$ mice. **B.** qPCR analysis of *Smarca4* mRNA level was performed in organoids treated with control or sg-*Smarca4* lentivirus. **C.** Representative images of organoids formed by crypts from *Setd2*$^{\text{ISC-WT}}$, and *Setd2*$^{\text{ISC-KO}}$ mice in the presence of control or sg-*Smarca4* lentivirus. Scale bars, 100 μm. **D-E.** Expression of stem cell markers (**D**), cell senescence-related genes (**E**) and FAO-related genes (**F**) in organoids in **Figure 6H**. For qRT-PCR, GAPDH was used as an internal control. The statistical data represent mean ± s.d. (n = 3 mice per genotype). Student's t-test: ns, non-sense. *P < 0.05. **P < 0.01. ***P < 0.001. All images are representative of n = 3 mice per genotype.
